# Soluble antigen arrays improve the efficacy and safety of peptide-based tolerogenic immunotherapy

**DOI:** 10.1101/2023.05.05.539161

**Authors:** Rebuma Firdessa-Fite, Stephanie N. Johnson, Martin A. Leon, Joshua O. Sestak, Cory Berkland, Remi J. Creusot

**Affiliations:** Columbia Center for Translational Immunology, Department of Medicine and Naomi Berrie Diabetes Center, Columbia University Medical Center, 650 West 168^th^ St, New York, NY 10032; Department of Pharmaceutical Chemistry, University of Kansas, 2095 Constant Avenue, Lawrence, KS 66047; Department of Chemistry, University of Kansas, 1251 Wescoe Hall Drive, Lawrence, KS 66045; Department of Chemical and Petroleum Engineering, University of Kansas,1530 West 15^th^ Street, Lawrence, KS 66045

**Author notes:** Corresponding author: Rémi J. Creusot., **Email:**.

**Keywords:** Soluble antigen arrays, modification and delivery of peptide, immunotherapy, autoimmune diabetes

## Abstract

Autoantigen-specific immunotherapy using peptides offers a more targeted approach to treat autoimmune diseases, but the limited *in vivo* stability and uptake of peptides impedes clinical implementation. We previously showed that multivalent delivery of peptides as soluble antigen arrays (SAgAs) efficiently protects against spontaneous autoimmune diabetes in the non-obese diabetic (NOD) mouse model. Here, we compared the efficacy, safety, and mechanisms of action of SAgAs versus free peptides. SAgAs, but not their corresponding free peptides at equivalent doses, efficiently prevented the development of diabetes. SAgAs increased the frequency of regulatory T cells among peptide-specific T cells or induce their anergy/exhaustion or deletion, depending on the type of SAgA (hydrolysable (hSAgA) and non-hydrolysable ‘click’ SAgA (cSAgA)) and duration of treatment, whereas their corresponding free peptides induced a more effector phenotype following delayed clonal expansion. Moreover, the N-terminal modification of peptides with aminooxy or alkyne linkers, which was needed for grafting onto hyaluronic acid to make hSAgA or cSAgA variants, respectively, influenced their stimulatory potency and safety, with alkyne-functionalized peptides being more potent and less anaphylactogenic than aminooxy-functionalized peptides. Both SAgA variants significantly delayed anaphylaxis compared to their respective free peptides. The anaphylaxis, which occurred in NOD mice but not in C57BL/6 mice, was dose-dependent but did not correlate with the production of IgG1 or IgE against the peptides. We provide evidence that SAgAs significantly improve the efficacy and safety of peptide-based immunotherapy.

**SIGNIFICANCE STATEMENT:** Peptide-based immunotherapy has several advantages over using full antigen as they are easy to synthetize, chemically modify and customize for precision medicine. However, their use in the clinic has been limited by issues of membrane impermeability, poor stability and potency *in vivo*, and in some cases, hypersensitivity reactions. Here, we provide evidence that soluble antigen arrays and alkyne-functionalization of peptides could be used as strategies to improve the safety and efficacy of peptide-based immunotherapy for autoimmune diseases by influencing the nature and dynamics of immune responses induced by the peptides.

## INTRODUCTION

Type 1 diabetes (T1D) is an autoimmune disease mediated by lymphocytes reactive to insulin-producing pancreatic β-cell antigens that result in insulitis and loss of β-cells. T1D is affecting millions of Americans and has currently no cure. Its control requires managing blood glucose levels with regular insulin administrations, which is cumbersome, does not prevent the development of long-term complications in many patients, and does not tackle the root cause of the disease. There is also an unmet need for safe therapies that can be applied early enough so that β-cells can be preserved and life-long dependence on exogenous insulin averted. To that end, antigen-specific immunotherapy (ASIT) offers a more targeted and selective way of disabling disease-specific autoreactive lymphocytes to treat T1D without dampening the whole immune system.

Using peptides as antigens for ASIT has several advantages over using full proteins. Peptides are easier to manufacture, chemically modify, and customize for precision medicine (using a “mix & match” approach with peptides covering multiple antigens and restricted to specific human leukocyte antigen (HLA) haplotypes) (1-3). However, the use of peptides for clinical applications has been limited by their short half-life (due to enzymatic degradation), high dispersion resulting in low cellular uptake “per cell”, and poor potency *in vivo* (4). In some cases, repeated administration also leads to anaphylactic events (5-8). These limitations can be overcome by using an efficient nanodelivery platform and chemical modification of peptides. On one hand, a desirable delivery modality for peptides would increase their resistance to enzymatic degradation, facilitate their drainage to lymphoid tissues, and enhance their uptake, ideally resulting in more efficient and persistent antigen presentation *in vivo* (5, 9). Several delivery platforms including nanoparticles (10-13), nanofibers (14), cell penetrating peptides (9, 15), and soluble antigen arrays (SAgAs) (5, 16) have been shown to address some of the above limitations of peptides and to improve the efficacy and safety of ASIT. In some cases, anaphylaxis caused by peptides in free form could be averted by ensuring slow release (17) or non-free forms (18). On the other hand, peptide modifications such as PEGylation (19), side chain stapling (20), retro-inverso-D-amino acid peptides (21, 22), and lipidation (23, 24) can also improve the efficacy and safety of peptide-based ASIT. For instance, lipophilic modification of InsB9-23 or addition of RLGL to WE14 peptides at the N-terminus was shown to enhance antigen presentation and to induce antigen-specific immune tolerance in models of T1D (25, 26). Acidic residues can contribute to peptide-induced anaphylactic reactions, and amino acid additions or substitutions that neutralize these charges prevented these effects (6, 7). Likewise, modification of antibody contact residues within the peptide could prevent antibody recognition and overcome the risk of peptide-induced anaphylaxis (27). These examples support the feasibility of overcoming inherent limitations of peptide-based ASIT using suitable delivery modalities and chemical modifications.

We recently used SAgAs as a multivalent, versatile, and effective peptide delivery modality that features multiple copies of antigenic peptides bound to hyaluronic acid (HA). SAgAs have advantageous properties such as a small size (less than 10 nm radius) and improved solubility (relative to free peptides) that allow their efficient delivery to lymphoid tissues. SAgAs enhance peptide uptake and persistence of antigen presentation (5, 16, 28, 29). Thus, SAgAs address many of the aforementioned limitations of peptide-based ASIT. Peptides used to produce SAgAs require N-terminal aminooxy (ao) or homopropargyl (hp) modification for conjugation of multiple copies to HA using aminooxy chemistry (producing hydrolyzable SAgAs (hSAgAs)) or using “click” chemistry (producing click SAgAs (cSAgAs)), respectively (5, 16, 30, 31). Following uptake of SAgAs by antigen-presenting cells (APCs), the peptides are released from the HA backbone (by hydrolysis at low pH for hSAgAs, or by mechanisms so far unclear but possibly involving degradation of HA itself for cSAgAs) and loaded onto MHC-II for presentation to CD4+ T cells. We previously showed that SAgAs carrying p79 mimotopes and 2.5 hybrid insulin peptides (2.5HIP) restore immune tolerance to β-cell antigens in non-obese diabetic (NOD) mice, protecting them from developing autoimmune diabetes in part by inducing regulatory CD4+ T cell populations (5, 30). The NOD mouse constitutes a polygenic and spontaneous model of autoimmune diabetes that shares many commonalities with human T1D (32, 33). Herein, we compared the safety and efficacy of hSAgAs and cSAgAs to their corresponding free peptides at equivalent peptide doses and characterized the dynamics and phenotypes of responding T cells.

## RESULTS

### Peptide modifications and dosing influence the dynamic of anaphylactic responses

Our previous studies (5, 30) focused on comparing single SAgAs (carrying p79 or 2.5HIP) versus a mix thereof (SAgA_mix_) at different doses. We reported that both peptides (as SAgA_mix_ at 2.5 nmol dose) were needed to achieve significant protection from disease (5). However, some mice treated with hSAgA_mix_ developed anaphylaxis over time following repeated dosing, which precluded a full long-term assessment of efficacy. Thus, we set out to determine the conditions that influence the incidence of anaphylaxis. At 0.5 nmol, hSAgA_mix_ treatment was safe, whereas it induced anaphylaxis after 8-11 weekly doses at 2.5 nmol (Fig.1A,B). The same dose given as free ao-peptide_mix_ (25 nmol peptide ≈ 2.5 nmol SAgA as SAgA carry ∼10 peptides in average) or higher dose of hSAgA_mix_ (12.5 nmol) greatly accelerated the incidence of anaphylaxis (Fig.1A,B). We then compared hSAgA_mix_ and cSAgA_mix_ (2.5 nmol) against the corresponding peptides used to produce them (ao and hp; 25 nmol). While cSAgA_mix_ was much safer than hSAgA_mix_ (5), the hp-modified peptide_mix_ was surprisingly also significantly less anaphylactogenic than its ao-modified counterpart (Fig.1C). Importantly, both SAgA forms were safer than their respective free peptides in delaying anaphylaxis (Fig.1A,C), underlining the safety advantage of SAgAs.

**Figure 1.**
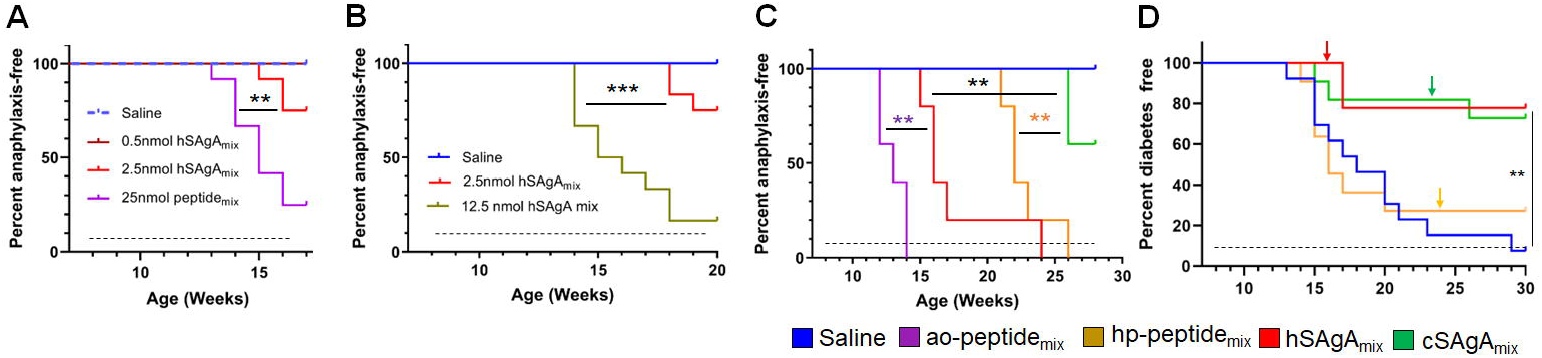
Incidence of anaphylaxis and diabetes in NOD mice. SAgAs mitigate anaphylaxis caused by free peptides in dose- and formulation-dependent manner (**A-C**). (**A**) Mice were treated with different doses of hSAgA_mix_ (n=12), ao-peptide_mix_ (n=12), or saline (n=12). (**B**) Incidence of anaphylaxis in mice treated with saline (n=12), 2.5 nmol (n=12), or 12.5 nmol (n=12) dose of hSAgA_mix_. (**C**) Comparison of incidence of anaphylaxis in mice treated with saline (n=5), hSAgA_mix_, and cSAgA_mix_ (2.5 nmol, n=5 mice/group) and their respective peptide_mix_ (ao and hp; 25 nmol, n=5 mice/group). (**D**) Incidence of diabetes in mice treated with saline (n=13), hp-peptide_mix_ (25 nmol, n=13), hSAgA_mix_, or cSAgA_mix_ (2.5 nmol, n=13). Arrows indicate tapering of the dose to 5 nmol each (peptide_mix_) or 0.5 nmol each (SAgA_mix_). In all panels, the dashed lines indicate the treatment period (weekly injections). Statistical analysis was performed using log-rank test for all panels.

### SAgAs, but not free peptides, efficiently block the development of autoimmune diabetes

In our previous study (5), the efficacy of the free peptides was not evaluated, nor was a side-by-side comparison of hSAgA_mix_ versus cSAgA_mix_ conducted. Given that the safe 0.5 nmol dose of SAgA was not significantly protective when given from beginning (5), we assessed the therapeutic efficacy of SAgAs and peptides in preventing diabetes in NOD mice using the 2.5 nmol dose but tapering to 0.5 nmol at the first sign of (or prior to) anaphylaxis (Fig.1D). The time of tapering (indicated by arrows) was consistent with the incidence of anaphylaxis assessed separately (Fig.1C), and we tapered cSAgA_mix_ at the same time as its free peptide equivalent for better comparison, even though no sign of anaphylaxis was observed at that time. No anaphylaxis was observed in any group following dose tapering. Our results indicate that (1) these peptides (p79 and 2.5HIP) in free form do not confer any protection (evident before dose tapering), (2) hSAgA and cSAgA have a comparable efficacy if anaphylaxis is averted, and (3) dose tapering does not result in substantial loss of protection, but it efficiently overcame anaphylaxis. Of note, we excluded ao-modified peptides in this study due to their high anaphylactogenic nature. These data underline the therapeutic advantage of SAgAs as a peptide delivery modality.

### The nature and dynamics of antigen-specific T cell responses depend on the mode of delivery and dosing period

Using adoptive transfer of antigen-specific T cells, we previously showed that SAgAs were more stimulatory than their corresponding free peptides at the same dose *in vivo*, inducing higher expression of anergy markers and more IL-10 than IFN-γ production, which was mostly contributed by the response to the p79 mimotope (5). We sought to investigate additional aspects of the T cell response for further insights into the mechanism of action. Using the same model and following a single dose of cSAgA_mix_ or hSAgA_mix_ (0.5 nmol each) or free soluble ao-peptide_mix_ (5 nmol each), the *ex vivo* recall response of transferred BDC2.5 CD4+ T cells was assessed 3 days after treatment by restimulating splenocytes with the ao-p79 or ao-2.5HIP peptides at 5 nM or their mix at 2.5 nM each. Analysis of culture supernatants 3 days later revealed the presence of substantial levels of Th2 cytokines (IL-4, IL-5, IL-10, IL-13), some Th1 cytokines (IFN-γ, TNF-α), as well as IL-6 and IL-22 induced by SAgAs (hSAgA_mix_ most prominently), but not by the free peptides (Fig.S1). Cytokine production was consistently and significantly higher with hSAgA_mix_ treatment than with cSAgA_mix_, except for IL-22, where cSAgA_mix_ induced the same or higher levels (Fig.S1). When a five-times lower dose was used (0.1 nmol SAgAmix, 1 nmol free peptides) under the same conditions, no cytokine response was detected upon peptide recall *ex vivo* (data not shown).

T cell responses are expected to evolve over the course of repeated antigen administrations and continuous exposure to the delivered epitopes. When assessing the response of endogenous T cells with MHC tetramers, cSAgA_mix_ induced a higher frequency of CD73+, CD73+ FR4+ (anergic cells), PD-1+, IL-10+, IL-10+ IFN-γ + and IL-2+ T cells among p79-reactive T cells than hSAgA_mix_, after two doses (three days apart) (Fig.S2,S3), 3 weekly doses, and 6 weekly doses (5). In contrast, after 23 weeks of continuous weekly dosing, these endogenous T cell responses looked dramatically different. By that time (end of the experiment shown in Fig.1D), the frequency of p79-reactive CD4+ T cells in cSAgA_mix_-treated mice had returned to levels close to the control group, significantly lower than in hSAgA_mix_-treated mice (Fig.2A), possibly as a result of deletion or contraction. Surprisingly, these p79-reactive T cells in hp-peptide_mix_-treated mice reached their highest frequency, indicating that T cells expand more slowly in response to free peptide than to SAgA. The high expression of CD73, FR4 and PD-1 seen at early time points was no longer observed, except to a limited extent in hSAgA_mix_-treated mice (Fig.2B-D). While p79-reactive T cells in cSAgA_mix_-treated mice were found at the lowest frequency out of all antigen-treated groups, these T cells retained the highest frequency of IL-10+ and IL-10+ IFN-γ+ (Fig.2E,F), Foxp3+ (Fig.2G and Fig.S4A), TIGIT+ and IL-2+ IL-10+ T cells (Fig.S4B,C). However, expression of IL-2+, TNF-α+ and IL-2+ TNF-α+ on p79-reactive T cells was lower in SAgA_mix_-treated groups than in hp-peptide_mix_-treated mice (Fig.S4D-F). IFN-γ+ (IL-10-) p79-reactive T cells did not change in frequency and were not significantly upregulated by the treatments at both early and late time points (Fig.S3D,S4G). Finally, expression of Lag3 and KLRG1 followed yet another pattern, with significant expression in mice treated with hp-peptide_mix_ and its SAgA equivalent (cSAgA_mix_), but not with hSAgA_mix_ (Fig.S4H,I).

**Figure 2.**
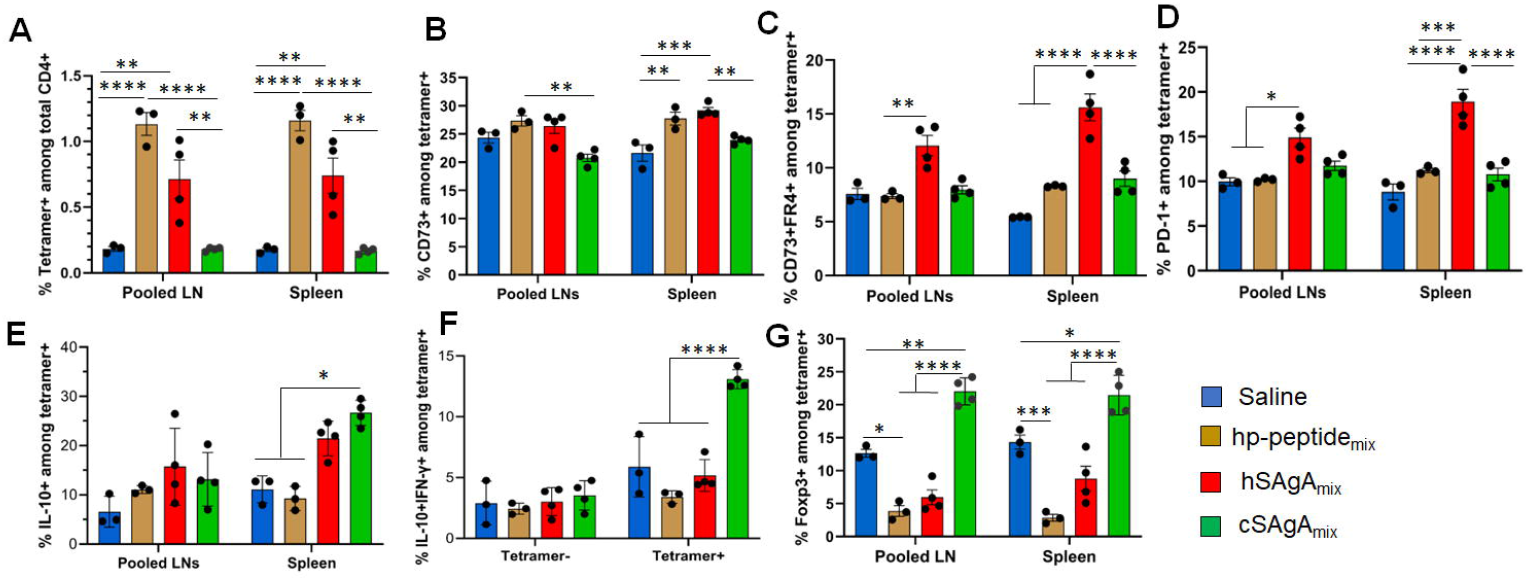
Comparison of polyclonal p79-reactive T cell responses between different modes of peptide delivery. T cell responses were measured in pooled lymph nodes (LNs) and spleen by flow cytometry after prolonged (23-week) weekly treatment with 2.5 nmol of hSAgA_mix_ or cSAgA_mix_, or 25 nmol of hp-peptide_mix_ (dose is for each peptide or SAgA (p79 and 2.5HIP) in the mix). Analysis was done on p79-tetramer+ CD4+ T cells and data are presented as: (**A**) % p79-tetramer+ cells among total CD4+ T cells, (**B**) % CD73+, (**C**) % CD73+/FR4+, (**D**) % PD-1+, (**E**) % IL-10+, (**F**) **%** IL-10+ IFN-γ+, and **(G)** % Foxp3+ among p79-tetramer+ CD4+ T cells. Data show the mean ± SEM from 3-4 mice per group. Statistical analysis was performed using two-way ANOVA/Tukey for all panels.

### Targets and mechanisms of the anaphylactic reaction to peptides

We also investigated why the different peptide delivery modalities led to very different kinetics of anaphylactic reactions (Fig.1C) and the mechanisms underlying these responses. We initially encountered this adverse effect when treating with hSAgA_mix_. Thus, to assess the target and type of anaphylactic response in an amplified and accelerated manner, we treated NOD mice with 10 nmol hSAgA_mix_ (a dose at which most mice developed anaphylaxis after only 5 weekly injections). Complete blood counts performed at the fifth dose indicate a dramatic elevation of all major leukocyte populations in the blood (Fig.3A). Serum collected at t = 0, 2 and 4 weeks was analyzed by multiplex assays to determine the level of different immunoglobulin isotypes. While levels of total IgG2a, IgG2b, IgG3 and IgM remained unchanged, IgG1 levels were significantly elevated after only two doses, and IgE levels significantly increased after four doses (Fig.3B). To assess the specificity of these antibodies, we performed ELISA using adsorbed ao-p79 or ao-2.5HIP peptides, followed by detection of specific IgG1 or IgE. Detection of anti-p79 IgG1 was evident at high serum dilutions, whereas that of anti-2.5HIP IgG1 required more concentrated serum (about 25x) (Fig.3C). However, low anti-2.5HIP IgG1 levels were highly variable and appear to preexist naturally as they were also detected in the untreated control group (Fig.3C), which is consistent with the fact that 2.5HIP is a naturally occurring epitope while p79 is an artificial mimotope. Interestingly, the later IgE response was directed at neither p79 nor 2.5HIP (Fig.3D). These data initially suggested that the anaphylactic response may be driven by an anti-p79 IgG1 response, which may have been facilitated by an early anti-p79 Th2 response to hSAgA_mix_ (Fig.S1). After subsequently observing that the free peptides led to even more severe and accelerated anaphylaxis, we repeated these studies to include all the treatment groups (Fig.1C) and using the same therapeutic dose as in preclinical studies (2.5 nmol SAgA or 25 nmol free peptide). Unexpectedly, both hSAgA_mix_ and cSAgA_mix_ led to significantly higher total IgG1 and IgE levels than the free peptides (ao- and hp-peptide_mix_) after dosing for 6 weeks (Fig.3E-H), whereas total IgG2a, IgG2b, IgG3 and IgM levels remained unchanged as before (data not shown). Moreover, both hSAgA_mix_ and cSAgA_mix_ induced higher levels of anti-p79 and anti-2.5HIP IgG1, and anti-p79 IgG2a than their corresponding free peptides (though not significantly, likely due to the lower dose used relative to the previous study) (Fig.3I-L and Fig.S5A,B). As before, no peptide-specific IgE were observed in all groups while peptide-specific IgG2b, IgG3 and IgM were not measured. Fewer time points could be assessed for hSAgA_mix_ and ao-peptide_mix_, due to the faster onset of anaphylaxis. These data indicate that the levels of anti-p79 IgG1 alone neither predict nor explain the occurrence of anaphylaxis in NOD mice. The levels of anti-p79 IgG2a were orders of magnitude lower than that of IgG1 in all groups (because detection required much less diluted samples) and were likely not playing a role in modulating the effects of IgG1.

**Figure 3.**
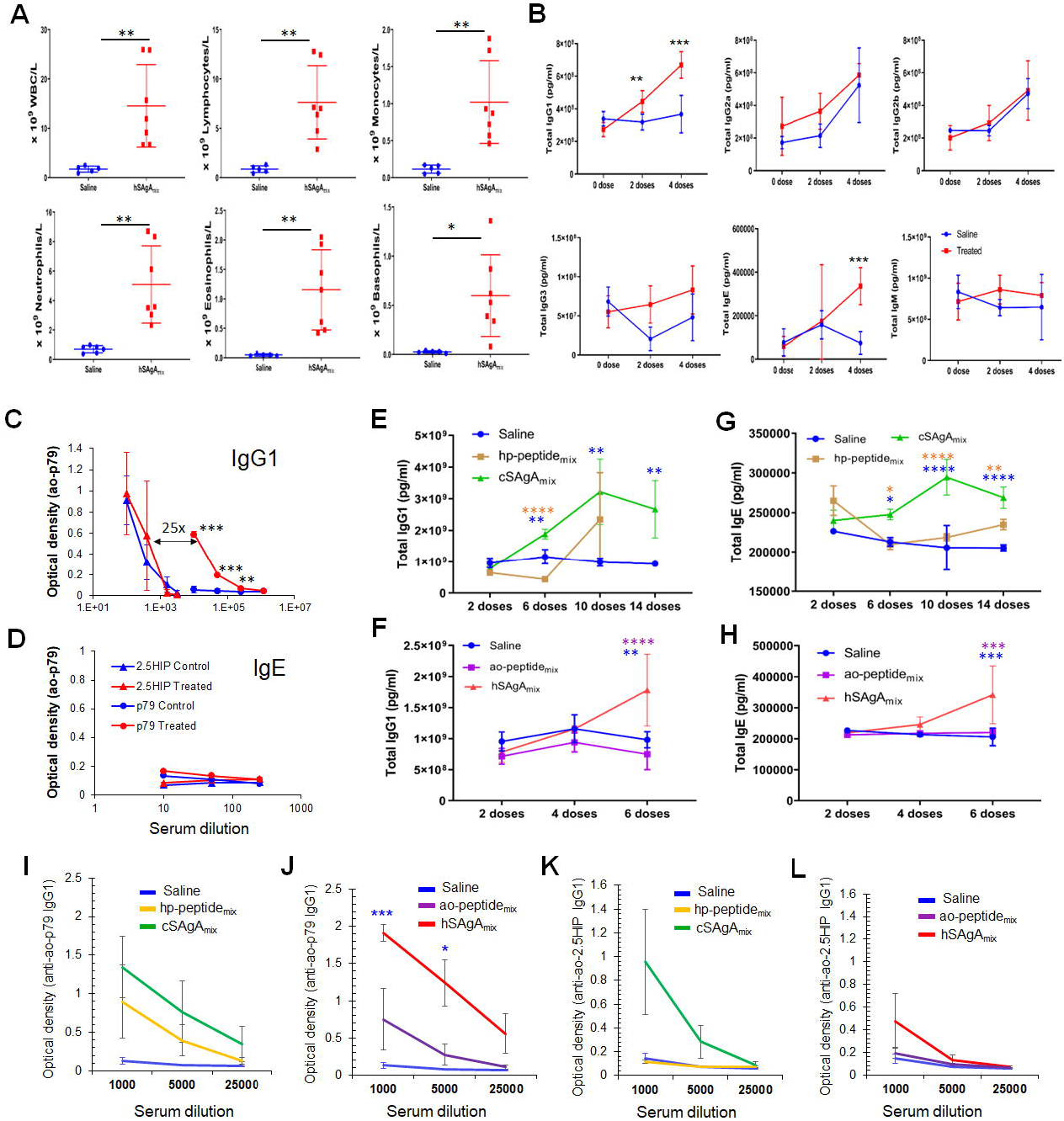
Characterization of the anaphylactic reaction to delivered peptides. (**A**) Changes in immune cell populations in the blood of mice developing anaphylaxis after 4 weekly doses of hSAgA_mix_ at 10 nmol each. (**B**) Levels of total IgG1, IgG2a, IgG2b, IgG3, IgE and IgM at baseline and after 2 and 4 weekly doses of hSAgA_mix_ (10 nmol each). No IgA was detected in all groups. Data in (A,B) show the mean ± SD from n=5 mice (control) and n=7 mice (treated). (**C**,**D**) Relative levels of IgG1 (C) and IgE (D) specific for 2.5HIP or p79 peptides with or without treatment with hSAgA_mix_ (4 weekly doses at 10 nmol each). Data in (C) show the mean ± SD from n=3 mice per group. (**E-H**) Comparison of total antibody isotype levels induced by both forms of SAgA and their corresponding peptides using the therapeutic dose of 2.5 nmol SAgA or 25 nmol free peptide as in preclinical studies and following the indicated number of doses. Total IgG1 (E,F) and IgE (G,H) were measured at 50,000 fold serum dilution. Other isotypes (IgG2a, IgG2b, IgG3 and IgM) were not changed at these serum dilutions. Data from cSAgAs/hp-peptides (E,G) and hSAgAs/ao-peptides (F,H) were separated for clarity and because of different kinetics of anaphylaxis development. Data show the mean ± SEM of 4-5 biological replicates. (**I-L**) Anti-p79 IgG1 (I,J) and anti-2.5HIP IgG1 (K,L) levels were measured at the indicated serum dilutions following treatment with saline, hp-peptide_mix_ or cSAgA_mix_ (10 doses) or ao-peptide_mix_ or hSAgA_mix_ (6 doses). Data from cSAgAs/hp-peptides (I,K) and hSAgAs/ao-peptides (J,L) were separated for clarity. Data show the mean ± SEM from 4-5 biological replicates. Statistical analysis was performed using non-parametric T-test for panels A,C,D,I-L, multiple T-test for panel B and two-way ANOVA/Tukey for panels E-H. The color of the star for significance level indicates the group with which the point immediately below the star is compared to.

Intriguingly, unlike treatment with hSAgA_mix_ at 2.5 nmol (each), treatment with a single hSAgA (p79 or 2.5HIP at 2.5 nmol) did not lead to anaphylaxis (Fig.S5C, (5)). In order to determine whether this difference was simply a dose issue, we tested the single hSAgA_p79_ at 5 nmol (a dose equal to both peptides of hSAgA_mix_ combined) and we observed that these mice had a comparable incidence of anaphylaxis as hSAgA_mix_-treated mice (Fig.4A), as well as comparable levels of total IgG1 (Fig.4B), IgE (Fig.4C), anti-p79 IgG1 (Fig.4D) and anti-p79 IgG2a (Fig.S5D) after 6 weekly doses. Thus, we identified a threshold between 2.5 nmol and 5 nmol at which both anti-p79 responses and anaphylaxis were enabled, and in hSAgA_mix_, the 2.5HIP peptide helped p79 (only at 2.5 nmol) reach that threshold.

**Figure 4:**
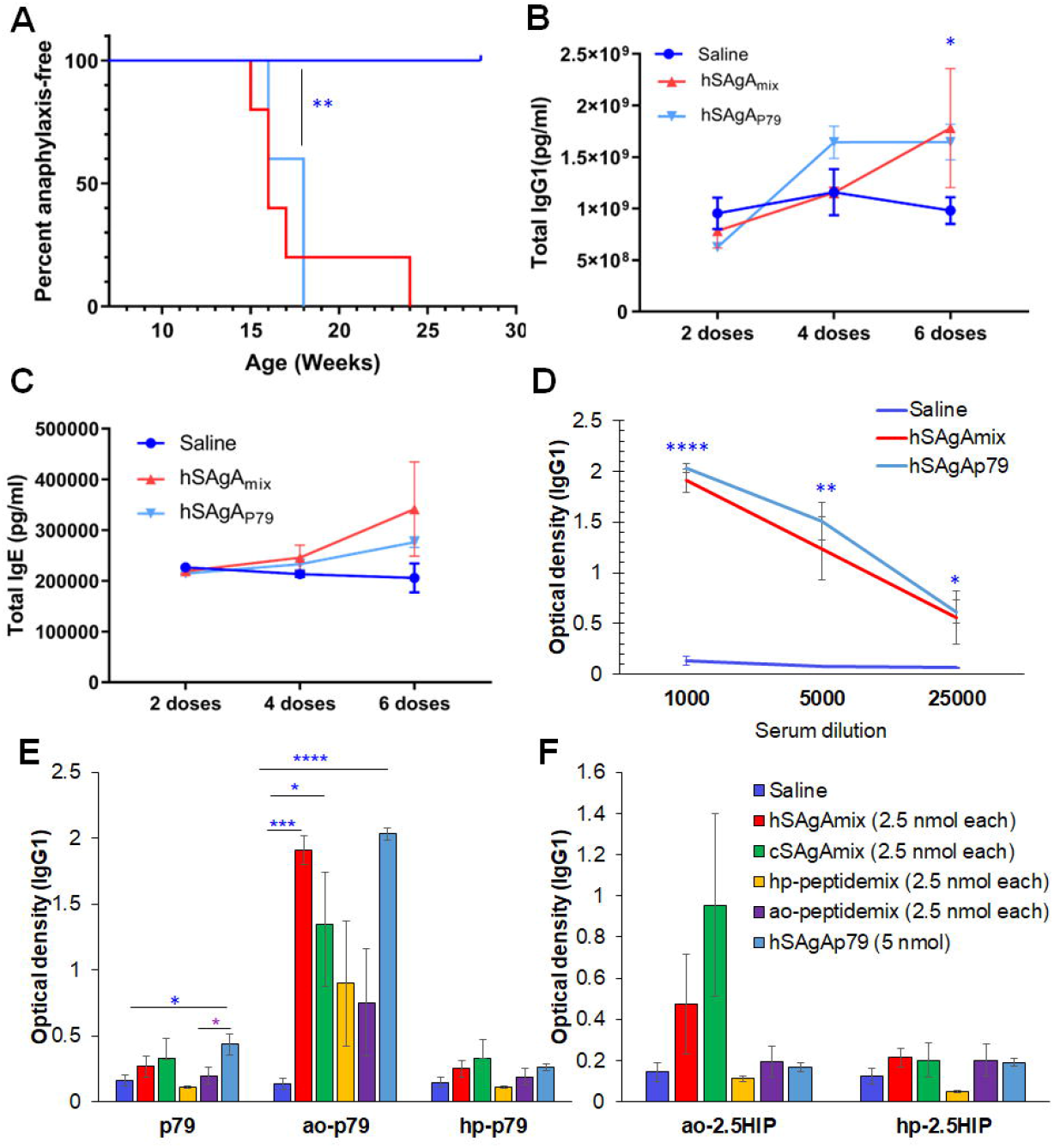
Comparison of anaphylaxis incidence and peptide-specific IgG1 levels between hSAgA_p79_ and hSAgA_mix_. (**A**) Incidence of anaphylaxis induced by hSAgA_mix_ at 2.5 nmol (each) versus single hSAgA_p79_ at 5 nmol (n=5 for each group). (**B-D**) Total IgG1 (B), total IgE (C) and anti-ao-p79 IgG1 (D) induced by single hSAgA_p79_ at 5 nmol as compared to hSAgA_mix_ at 2.5 nmol (each). (**E-F**) Peptide-specific IgG1 reactivity against ao-modified, hp-modified and unmodified p79 (E) and against ao-modified or hp-modified 2.5HIP (F) after 6 weekly doses for hSAgA_mix_, hSAgA_p79_, and ao-peptide_mix_ and after 14 weekly doses for cSAgA_mix_ and hp-peptide_mix_ at 1:1000 serum dilution. Data in (B-F) show the mean ± SEM from 4-5 biological replicates. Statistical analysis was performed using two-way ANOVA/Tukey for panels B-C while non-parametric T-test was applied for panels D-F. The blue stars indicate the level of significance when comparing single hSAgA_p79_ to saline control.

Because ao-peptide_mix_ and its corresponding hSAgA_mix_ were the most anaphylactogenic in our treatments, we considered the possibility of a response against the modified portion of the peptide. Thus, p79, ao-p79, hp-p79, ao-2.5HIP and hp-2.5HIP peptides were all assessed as targets in our ELISA assays to detect antigen-specific IgG1, IgG2a, and IgE levels induced by the different treatments. Surprisingly, both anti-p79 and anti-2.5HIP IgG1 levels were more pronounced against the adsorbed ao-modified peptides than the hp-modified peptides or unmodified peptides, regardless of the treatment (Fig.4E,F), while no peptide-specific IgE was detected for all conditions. Peptide modifications also accounted for differences in stimulatory activity: hp-modified p79 was 100x more stimulatory for BDC2.5 T cells than unmodified and ao-modified p79 *in vitro* ((30),Fig.S6A,B). Likewise, hp-modified 2.5HIP was 10x more stimulatory than ao-modified 2.5HIP (Fig.S6C-F).

### The anaphylactic reaction to delivered peptides is strain-dependent

Higher frequency of unwanted allergic responses has been associated with autoimmune diseases (34-36), thus we also evaluated our most anaphylactogenic mixes (hSAgA_mix_ at 2.5 nmol and ao-peptide_mix_ at 25 nmol) in the C57BL/6J (B6) mouse strain, which is not prone to develop autoimmune disease spontaneously. We continuously treated B6 mice with 19 weekly injections, past the time when anaphylaxis had occurred in all NOD mice, and none of them developed any sign of anaphylaxis (Fig.5A). For the last treatment, these mice received a higher dose of 10 nmol hSAgA_mix_ or 100 nmol ao-peptide_mix_, and still did not show any sign of anaphylaxis. Separately, we tested whether discontinuing the treatment for a prolonged period before resuming may overcome the development of anaphylaxis in NOD mice. NOD mice (n=16) were treated with hSAgA_**mix**_ at 2.5 nmol with 9 weekly doses until some of them started developing anaphylaxis (6/16). After 5 or 15 weeks of treatment interruption, the NOD mice were rechallenged with a single injection of hSAgA_**mix**_ at 2.5 nmol and all the mice that received the challenge (n=5 per interruption period) immediately developed fatal anaphylaxis (Fig.5B). Overall, treatment interruption did not appear to reset or delay the development of anaphylaxis, and once the NOD mice were sensitized with 5-9 weekly treatments with hSAgA_mix_ or free ao-peptide_mix_, they remained so for a prolonged period, whereas B6 mice were completely resistant to even continuous treatment at equivalent dose.

**Figure 5.**
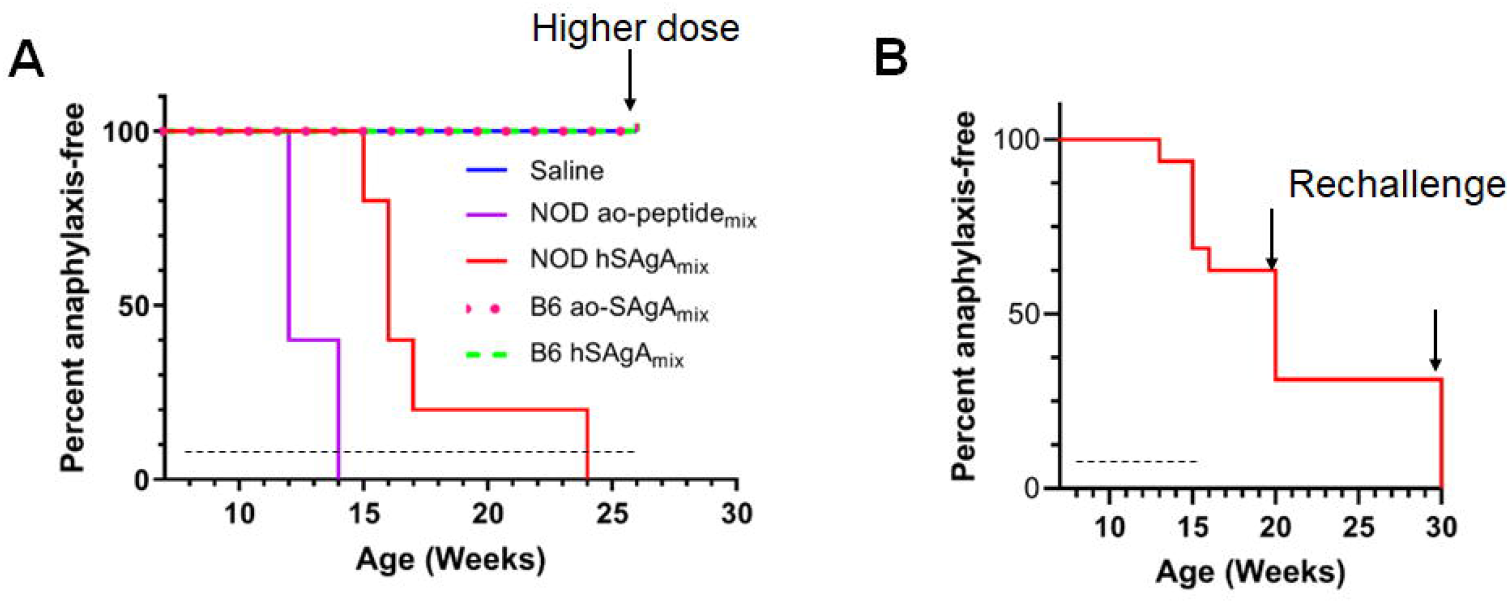
Incidence of anaphylactic reaction to ao-peptide_mix_ and hSAgA_mix_ is strain-dependent. (**A**) Incidence of anaphylaxis in NOD vs B6 mice (n=5 per strain) treated weekly with a 2.5 nmol dose of hSAgA_mix_ or 25 nmol of the ao-peptide_mix_ or saline. B6 mice also received an injection of a higher dose of 10 nmol hSAgA_mix_ or 100 nmol ao-peptide_mix_ as the final treatment. (**B**) Incidence of anaphylaxis in NOD mice in which treatment was resumed after a prolonged pause. Mice were initially treated with 2.5 nmol hSAgA_mix_ weekly (for 8 weeks), and treatment was discontinued when some mice (n=6) succumbed to anaphylaxis. Among the remaining 10 mice, 5 were treated again after 5 weeks with a single injection of hSAgA_mix_ at 2.5 nmol, and the other 5 were treated likewise after 15 weeks. In both cases, all mice developed fatal anaphylaxis. Dashed lines indicate the period of weekly treatments, arrows indicate a time point at which mice were rechallenged with a single injection of hSAgA_mix_ at 10 nmol (**A**) or 2.5 nmol (**B**).

## DISCUSSION

SAgAs constitute a versatile peptide delivery modality that overcomes many inherent limitations of peptide-based immunotherapy to induce immune tolerance through different mechanisms involving T cells and B cells (5, 16, 28-31, 37, 38). In this study, we showed that the efficacy and safety of peptide-based immunotherapy may depend on the use of desirable peptide modifications and/or the type of delivery modality. While both SAgA variants efficiently protected NOD mice from spontaneously developing T1D, their corresponding soluble peptides failed to protect the mice at equivalent doses, showing the critical role of the delivery modality to achieve therapeutic efficacy. Once immune tolerance is established by an initial high dose (2.5 nmol) of SAgAs given for several weeks, a later dose tapering (0.5 nmol) is sufficient to maintain protection and can completely abrogate the risk of anaphylaxis. As expected, the T cell dynamics following the treatment with the two SAgA variants and free soluble peptides were considerably different. Both SAgA variants induced more regulatory or anergic T cell phenotypes (CD73+ FR4+, PD-1+, IL-10+) and significantly less IL-2+ and TNF-α+ antigen-specific T cells than free peptides after prolonged treatment. In fact, the T cell responses induced by the latter, at a time of considerable clonal expansion, were more of an effector phenotype, with distinct upregulation of KLRG1, lack of anergy, exhaustion and regulation marker upregulation, unlike what was seen with SAgAs when T cells were expanding (5). Overall, cSAgAs induced stronger initial immune responses than hSAgA variants or their free peptides, and its continued weekly dosing may have resulted in conversion to regulatory T cells and/or deletion of autoreactive T cells, as evidenced by the low frequency of tetramer+ T cells, a higher proportion of which expressed Foxp3. hSAgAs induced weaker initial T cell responses than cSAgAs and their continued weekly dosing may have maintained anergic and/or IL-10+ T cells that protected the mice from T1D. In contrast, the free soluble peptides induced a weak response initially, but its continuous administration eventually resulted in increased frequency of antigen-specific T cells, which adopted a more classic effector phenotype and had no effect on disease progression in NOD mice. This weaker response to the free peptide *in vivo* was also evident from the cytokine analysis data with no detectable levels beyond the control at early time point following adoptive transfer of the transgenic BDC2.5 CD4+ T cells and *ex vivo* restimulation of splenocytes. On the contrary, recall response by splenocytes from mice treated with SAgA_mix_ (particularly hSAgA_mix_) resulted in significantly higher production of Th2 cytokines as compared to the free ao-peptide or saline control, which may have also contributed to the protection. Interestingly, Lag3 expression on p79-reactive T cells was prominent in response to both hp-peptide and cSAgAs, suggesting that the greater stimulatory capacity conferred by the hp modification may influence Lag3 induction. Thus, the phenotype of antigen-specific T cells after few administrations versus prolonged treatment differed substantially and was also dependent on the type of delivery modality used.

Modification of peptides was needed for SAgA formulations, and we unexpectedly found that the hp-modification improved both the potency and safety of the peptides. The hp-modified peptides were more stimulatory than their corresponding ao-modified peptides or unmodified p79 form at equivalent concentrations. In line with this, we previously showed that hp-p79 and ao-p79 secondary structures were different, possibly because of differences in the hydrophobicity of the modification (30). Thus, this secondary structure difference between the two modified peptides might alter the fitting and orientation of the peptides on the MHC groove for presentation to T cells. Whether this effect of hp modification on improving the peptide’s stimulatory potency could be generalizable to peptides restricted to other MHC II haplotypes/HLA alleles remains to be addressed. Moreover, hp-functionalized peptides and their SAgA form (cSAgA_mix_) significantly delayed the incidence of systemic anaphylaxis as compared to the ao-functionalized peptides and hSAgA_mix_, pointing to a role of hp-peptide modification in minimizing the development of anaphylaxis. The lower levels of Th2 cytokines induced by cSAgA_mix_ compared to hSAgA_mix_ may provide one explanation for the lower anaphylactogenic potential of cSAgA_mix_. Indeed, a Th2 cytokine profile has been associated with a risk of developing anaphylactic reactivity to therapeutic peptides although shifting the antigen-specific immune response towards Th2 is one mechanism by which several ASITs were shown to suppress the progression of autoimmunity in mice (36, 39-41).

Systemic anaphylaxis is rapid in onset and features potentially life-threatening immune reactions mediated by immunologic or non-immunologic causes (7, 27, 42). The mechanism responsible for most cases of anaphylaxis in humans involves the classic pathway which is mediated by IgE engagement with high-affinity Fc receptor (FcεRI) on mast cells and basophils, thereby inducing the release of inflammatory mediators (43). In humans, it may also be mediated by IgE and IgG1 combined (43, 44). For the alternative pathway, which is IgG-mediated and mainly reported in rodent models, platelet-activating factor (PAF), rather than histamine, is an important mediator in actively immunized mice and is released by basophils, monocytes/macrophages and neutrophils activated via their Fc gamma receptors (FcγRs) (44, 45). In this study, we report an antibody response that was consistently directed at p79 (an artificial mimotope), and in some cases at 2.5HIP (a natural β-cell neoepitope). Sudden exposure to large amounts of an artificial mimotope may explain stronger B cell reactivity, although anaphylactic reactions have been reported in NOD mice injected with natural islet peptides (46, 47). Paradoxically, the two SAgA variants were associated with elevated levels of total IgG1 and IgE, peptide-specific IgG1 and Th2 cytokines relative to the free peptides, but were less anaphylactogenic. This was surprising given that whole allergen and multivalent antigens are more likely, for example, to crosslink IgE and induce mast cell and basophil degranulation and to provide 3D conformations that are optimal to function as B cell epitopes compared to soluble peptides (48, 49). SAgAs are more stimulatory than free peptides at equivalent doses, in part by increasing peptide load per cell and, in turn, the avidity of antigen recognition by T cells. We postulated that SAgAs may consequently induce more Th1-associated peptide-specific IgG2a that would outcompete peptide-specific IgG1, thereby reducing their anaphylactogenic potential. However, both SAgAs and peptides induced much lower levels of peptide-specific IgG2a than IgG1, making it difficult to reconcile the high anaphylaxis incidence of free peptides with their reduced or absent induction of immunoglobulins and cytokine responses. Alternatively, the higher frequency of antigen-specific IL-10+ regulatory T cells induced by SAgAs may contribute to a better control of anaphylactic responses in the long-term (50, 51).

Unlike NOD mice, we found B6 mice to be completely resistant to anaphylaxis induced by hSAgA_mix_ or the ao-peptide_mix_ even after an extended dosing period and the use of 4 times higher dose for rechallenge. Consistent with our finding, NOD mice were shown to be more susceptible to anaphylactic shock than other common mouse strains (41). Likewise, repeated injections of human albumin or alpha-1 antitrypsin, but not the mouse versions, were associated with a high rate of fatal anaphylaxis in NOD mice (80-100%) while this occurrence was limited in NOR mice (10%) and absent in BALB/c and B6 mice, indicating a greater susceptibility of an autoimmune-prone strain (NOD mice) to hypersensitivity reactions to ASITs than other strains (52). While the resistance of B6 mice may be explained by an MHC class II haplotype (I-A^b^) potentially unable to present p79 and 2.5HIP, the reported difference between NOD and NOR mice (both with the same I-A^g7^ haplotype) (52) suggests that the ability to present antigens only partially explains susceptibility to anaphylaxis. Moreover, studies in the mouse model of multiple sclerosis indicated that anaphylactic reactions may be elicited with self-peptides that are not presented in the thymus, indicating that a lack of central tolerance, and possibly of thymic regulatory T cells, may create permissible conditions for anaphylaxis (36). Independent from antigen presentation, chemically modified peptides may directly bind to and activate mast cells (53), which could provide an explanation for the greater anaphylactogenic potential of the free peptides, as their binding would be hindered in SAgA form. However, this does not seem to be the case, as these peptides had no effect in B6 mice. Therefore, we propose that autoimmune strains/individuals may be more prone than others to developing anaphylactic reactions to peptides in ASIT, in part due to susceptible MHC alleles and impaired T cell regulation. As novel antigenic peptides are evaluated in ASIT, it is ineluctable that some may cause anaphylaxis, and thus, delivery via SAgA offers an additional safety advantage in mitigating this risk.

## MATERIALS AND METHODS

### Mice

All mice were used according to approved protocols by Columbia University Institutional Animal Care and Use Committee. Female NOD mice (Jax #001976) were obtained from The Jackson Laboratory at 7 weeks of age and were directly used for preclinical (treatment) and anaphylaxis incidence studies one week after their arrival in the animal barrier facility of the Columbia Center for Translational Immunology. For mechanistic studies, BDC2.5 T cell receptor (TCR) transgenic mice (Jax #004460) and NOD.CD45.2 congenic mice (Jax #014149) were originally procured from The Jackson Laboratory but bred together to produce BDC2.5 mice with the CD45.2 congenic marker and maintained in our animal barrier facility. They were used at 8-12 weeks of age as donors of antigen-specific T cells for *in vivo* tracking after adoptive transfer. Female NOD mice (8-12 weeks of age) were used for short mechanistic studies involving adoptive transfer (recipient mice) and MHC tetramer analysis. C57BL/6J mice (Jax #000664) were bred in our animal barrier facility and were used for anaphylaxis study as non-autoimmune-prone strains.

### Synthesis of hydrolysable hSAgA and non-hydrolysable (“click”) cSAgA

Alkyne-functionalized p79 bearing an N-terminal 4-pentynoic acid (homopropargyl) modification, alkyne-functionalized 2.5HIP bearing an N-terminal alkyne polyethylene glycol group modification, or aminooxy-functionalized p79 or 2.5HIP were purchased from PolyPeptide. SAgAs were synthesized by co-grafting approximately 10 hp-peptides to azide-functionalized HA to make cSAgAs using click chemistry or linking approximately 10 ao-peptides to HA to make hSAgAs using oxime conjugation chemistry and their physicochemical natures were characterized as previously reported (5, 16).

### Preclinical studies

At 8 weeks of age, NOD mice were treated subcutaneously (s.c.) at the neck fold with saline, SAgA_mix_, or peptide_mix_ weekly at various peptide doses, ranging from 5 nmol to 125 nmol for each peptide from the mixture. Their blood glucose was monitored weekly (up to 30 weeks of age), and mice were diagnosed as diabetic after two consecutive blood glucose levels greater than 250 mg/dL.

### Assessment of anaphylactic incidences and responses

Female NOD and C57BL/6J mice were treated weekly with hSAgA_p79_ (5 nmol), hSAgA_mix_ (2.5-12.5 nmol each), ao-peptide_mix_ (25-125 nmol each), cSAgA_mix_ (2.5 nmol each), hp-peptide_mix_ (25 nmol each), or saline for the period indicated in the figure legend. Incidence of anaphylaxis was recorded when mice developed fatal systemic anaphylaxis with typical type I hypersensitivity symptoms such as trouble breathing and loss of consciousness within 30 min. Blood samples were drawn every two weeks and were analyzed for complete blood count at the Department of Comparative Medicine using a Genesis instrument (Oxford Science Inc.). Serum samples were assessed for titers of different antibody isotypes using LegendPlex Mouse Immunoglobulin Isotyping Panel and IgE ELISA (BioLegend) according to manufacturer’s instructions. For antigen-specific indirect ELISA, the plate was coated with p79, ao-p79, hp-p79, ao-2.5HIP, or hp-2.5HIP (each at 10 μg/ml) for each immunoglobulin at 4ºC overnight. After the plates were washed and blocked, four serially diluted serum samples from mice treated with saline, hSAgA_p79_, hSAgA_mix_, cSAgA_mix_, ao-peptide_mix_, or hp-peptide_mix_ were added. The plates were washed and detection antibodies (biotinylated mouse anti-IgG1, anti-IgG2a or anti-IgE from BioLegend) were added and incubated at room temperature for 1 hour. Finally, avidin-horseradish peroxidase solution was added to each well for a 30-min incubation at room temperature. The absorbance was measured at 450 nm and 570 nm on the same day.

### T cell response analysis

To evaluate BDC2.5 CD4+ T cell responses induced by unmodified free peptides vs modified *in vitro*, splenocytes from NOD.BDC2.5.CD45.2 mice were labeled with VCPD (eBioscience) and co-cultured (2×10^5^ total cells/well) in the presence/absence of titrated p79, ao-p79, hp-p79, ao-2.5HIP, or hp-2.5HIP at 10-fold serial dilutions ranging from 10pM to 1μM concentrations. After 3 days of co-culture at 37°C and 5% CO_2_, the splenocytes were analyzed for activation markers (CD25) and proliferation by flow cytometry. For the mechanistic studies, female NOD mice were treated weekly by s.c. injection of saline or soluble peptide_mix_ (25 nmol per peptide), or hSAgA_mix_ (2.5 nmol each) or cSAgA_mix_ (2.5 nmol each) for the indicated period in the legend. Spleen and various lymph modes (LNs) including pancreatic LNs, pooled axillary and brachial LNs, or pooled LNs (pool of axillary, brachial, cervical, mesenteric and/or pancreatic LNs) were collected, and single cells suspensions were prepared. Analysis of polyclonal T cell responses by flow cytometry was performed at two time points: early time point (after two injections three days part) and late time point (following 23 weekly injections). Multiple panels were used to assess surface markers (CD4, CD25, CD44, CD73, FR4, Lag3, PD-1, KLRG1 and TIGIT), intracellular cytokines (IFNγ, TNF-α, IL-2, and IL-10) and Foxp3 staining. Intracellular cytokine staining was performed after a 4-hour incubation with PMA (0.1 μg/mL), ionomycin (40 μg/mL), brefeldin A (1.5 μg/ml) and monensin (1 μM) using CytoFix/CytoPerm kit (BD Bioscience). Likewise, intracellular staining for transcription factor (Foxp3) was performed on using True-Nuclear Factor kit (Biolegend) following manufacturer’s instructions. To identify endogenous antigen-specific T cells, I-A^g7^/p79 (AAAAVRPLWVRMEAA on APC) tetramer from the NIH Tetramer Core Facility was used. Fortessa (BD) was used for data acquisition by flow cytometry.

### Cytokine analysis

NOD mice were untreated or treated with a single dose of soluble ao-peptide_mix_, hSAgA_mix_ or cSAgA_mix_ at 1 nmol or 5 nmol of peptide doses for each peptide (p79 or 2.5HIP) in the mix via s.c. route. At the same time, the mice received 5×10^5^ purified and Violet Cell Proliferation Dye (eBioscience) labeled BDC2.5 CD4+ CD25-T cells (with CD45.2 congenic marker) by intravenous injection. Three days after the treatment, the spleen was isolated and splenocytes (3×10^5^/well) were cultured in the presence of ao-p79 (5 nM), ao-2.5HIP (5 nM), or their mix (2.5 nM each) to assess cytokine recall responses *ex vivo*. After four days of *ex vivo* culture, thirteen cytokines, namely IFN-γ, TNF-α, IL-6, IL-2, IL-4, IL-5, IL-9, IL-10, IL-13, IL-17F, IL-17A, IL-21, and IL-22 were measured in the culture supernatants using the LEGENDPlex Mouse T Helper Cytokine Panel kit (BioLegend) following manufacturer’s instructions.

### Statistical and Data Analysis

GraphPad Prism 4.0 was used to generate all graphs including the Kaplan-Meier curve and to perform statistical analyses. The log-rank (Mantel-Cox) test was used to calculate incidence diabetes and anaphylaxis. Two-way analysis of variance (ANOVA) with Sidak’s post-hoc test correction or Tukey’s multiple comparisons, and/or unpaired T-tests were performed in other studies as indicated in legends. Flow cytometry data were analyzed with FCS Express 7. The threshold for statistical significance was set to p<0.05 (* p<0.05, ** p<0.01, *** p<0.001, **** p<0.0001).

## Data availability

All study data are included in the article. Unprocessed data are available from the corresponding author upon reasonable request.

## Author contributions

RFF performed experiments, analyzed data, and wrote the first draft. RFF and RJC designed experiments. RJC and CB directed the research and edited the manuscript. SNJ, MAL, JOS and CB produced and characterized SAgAs. All authors approved the manuscript.

## ACKNOWLEDGMENTS

We thank Dr. Joshua Milner for a helpful discussion of the data. RFF was funded by postdoctoral fellowship 1-19-PMF-022 from the American Diabetes Association. SNJ and MAL were supported by the National Institutes of Health Graduate Training Program in Dynamic Aspects of Chemical Biology Grant (T32GM008545) from the National Institute of General Medical Sciences. These studies were originally funded by Juvenile Diabetes Research Foundation (JDRF) grant 2-SRA-2017-312-S-B to JOS. RFF and RJC were also supported by grant R01DK127778. Research reported in this publication was performed using the CCTI Flow Cytometry Core, supported in part by awards S10RR027050, S10OD020056 and P30DK063608. We thank the NIH Tetramer Core Facility, supported by contract HHSN272201300006C from the National Institute of Allergy and Infectious Diseases, for the MHC tetramer used in these studies.

The authors declare no conflict of interest pertaining to these studies.

