## Supplemental figures for "Soluble antigen arrays improve the efficacy and safety of peptide-based tolerogenic immunotherapy"

### Supplementary Figures

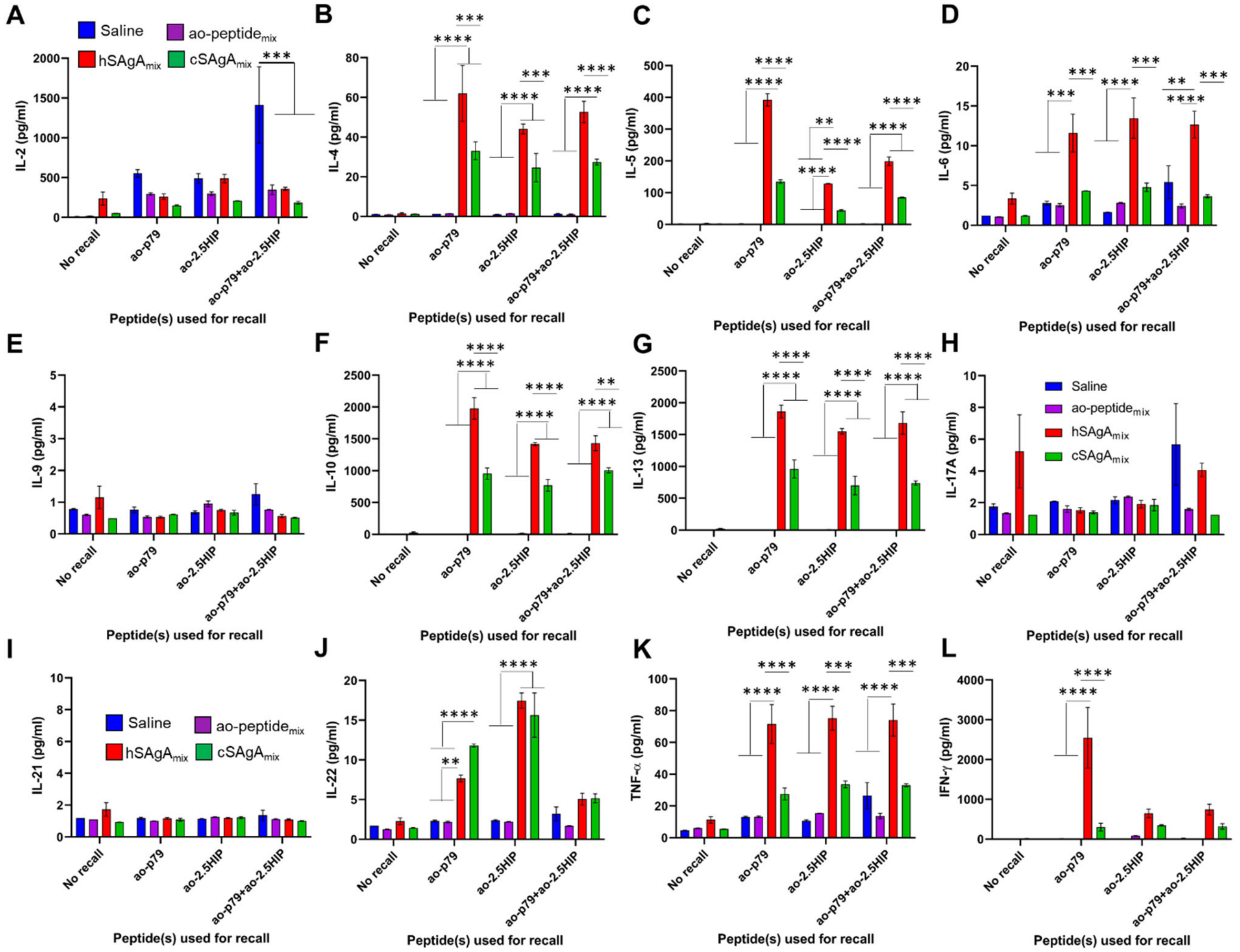


**Figure S1**. **Assessment of cytokines recall responses show production of substantial levels of Th2 cytokines induced by SAgAs as compared to free peptides**. NOD mice received 5x$10^5$ BDC2.5 CD4+ T cells adoptively and were treated with a single dose of cSAgA$_{mix}$ or hSAgA$_{mix}$ (0.5 nmol each) or free ao-peptide$_{mix}$ (5 nmol each). After 3 days, splenocytes were isolated and restimulated with ao-p79 or ao-2.5HIP peptides at 5 nM or their mix at 2.5 nM each. After another 3 days, culture supernatant was collected and assessed for IL-2 (**A**), IL-4 (**B**), IL-5 (**C**), IL-6 (**D**), IL-9 (**E**), IL-10 (**F**), IL-13 (**G**), IL-17A (**H**), IL-21 (**I**), IL-22 (**J**), TNF-α (**K**) and IFN-γ (**L**) levels using the LEGENDPlex Mouse Th1/Th2 Cytokine Panel kit (BioLegend) following manufacturer's instructions. Data show the mean ± SEM from 3 biological replicates. Statistical analysis was performed using two-way ANOVA/Tukey for all panels.

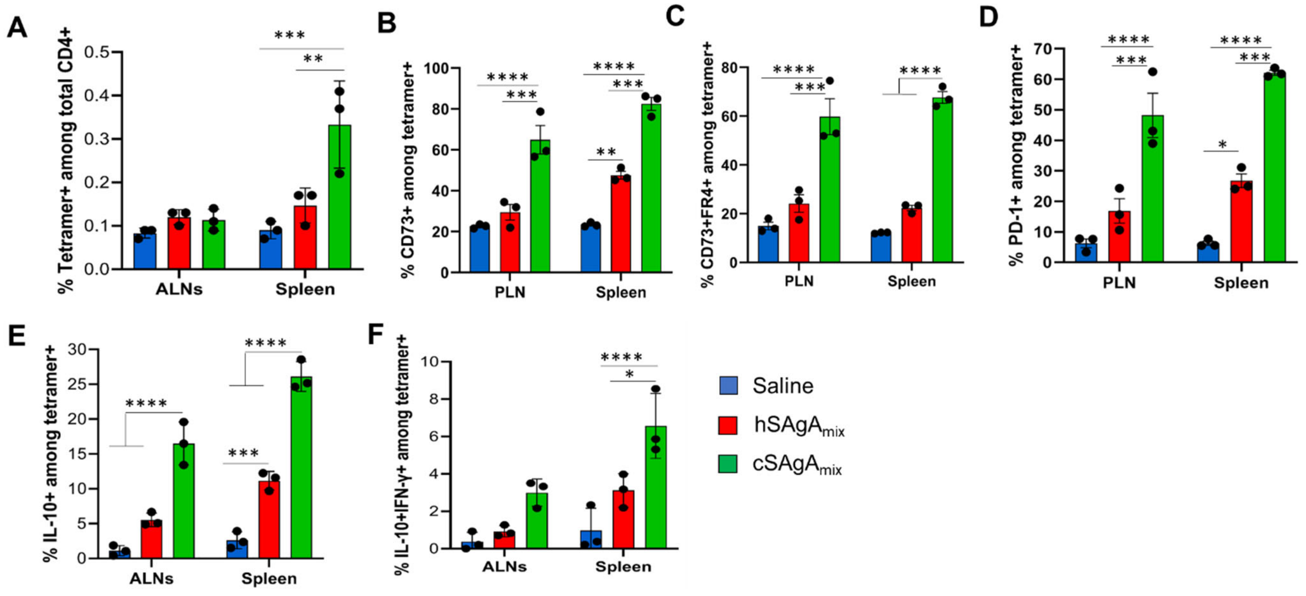


**Figure S2**. **Comparison of tolerance-associated markers induced on antigen-specific T cells by hSAgA$_{mix}$ vs cSAgA$_{mix}$ at early time point.** Phenotype of antigen-specific T cells induced by the two forms of SAgAs in lymph nodes and spleen: frequency of tetramer+ cells among total CD4+ T cells (**A**), and of CD73+ (**B**), CD73+ FR4+ (**C**), PD-1+ (**D**), IL-10+ (**E**) and IL-10+ IFN-γ+ (**F**) cells among p79-reactive T cells, after two doses (three days apart). Data show the mean ± SEM from 3 mice per group. Statistical analysis was performed using two-way ANOVA/Tukey for all panels. ALN is the pool of brachial and axillary lymph nodes. ALN was used for additional panels (intracellular staining) as there were not enough cells from PLNs for multiple panels.

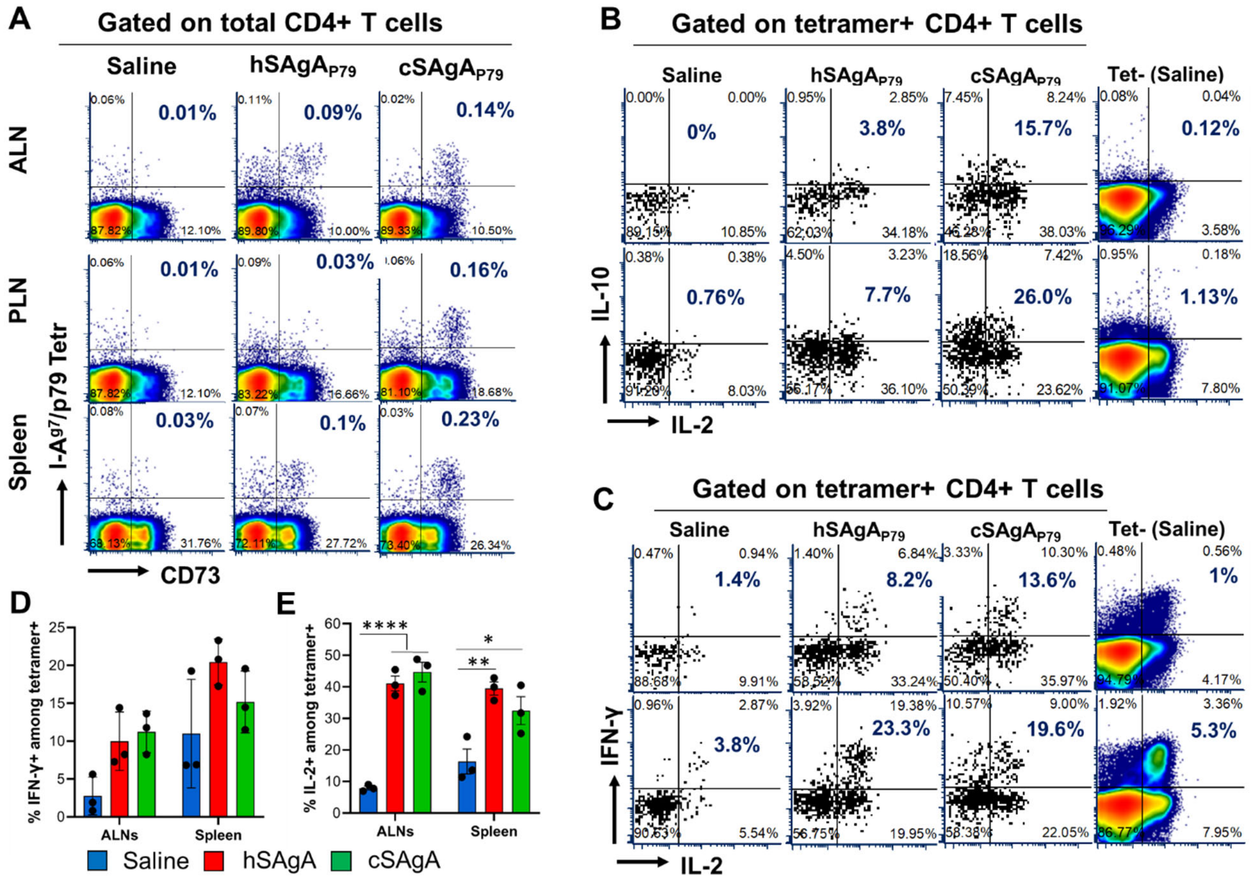


**Figure S3**. **Antigen-specific T cells responses induced by hSAgA$_{mix}$ versus cSAgA$_{mix}$ at an early time point.** Representative dot plots showing the percentage of tetramer+ CD73+ cells among total CD4+ T cells (**A**), IL-10 versus IL-2 expression (**B**), and IFN-γ versus IL-2 expression (**C-E**) cells among p79-reactive T cells than hSAgA$_{mix}$, measured 3 days after two treatments (three days apart). The percentage highlighted on each representative dot plot is for tetramer+ CD73+ among total CD4+ T cells (A), total IL-10+ (B) and total IFN-γ+ (C) cells among tetramer+ CD4+ T cells. The bar graphs show the mean ± SEM from 3 mice per group. Statistical analysis was performed using two-way ANOVA/Tukey for all panels. ALN is a pool of brachial and axillary lymph nodes.

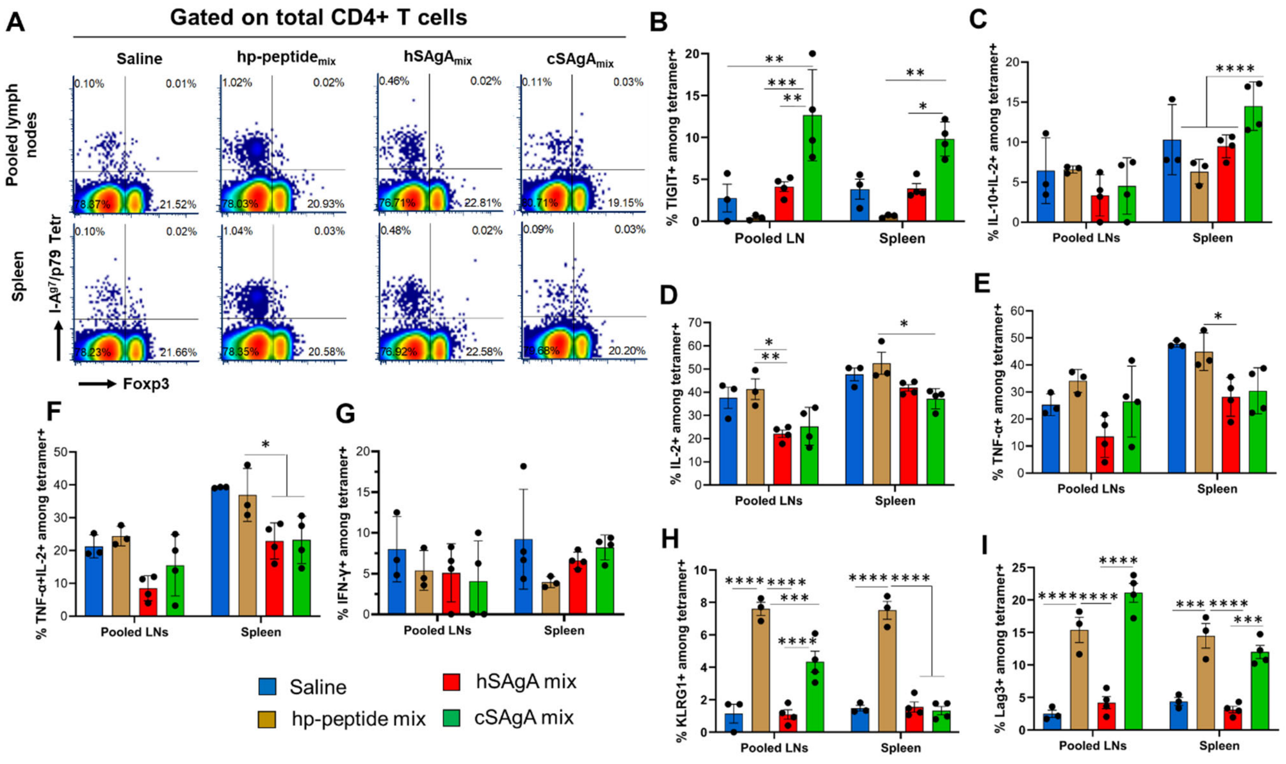


**Figure S4: Phenotype of antigen-specific T cells induced by SAgAs following prolonged weekly dosing.** (**A**) Representative dot plots of p79-reactive T cells gated on total CD4+ T cells 3 days after the last (23[rd]) weekly treatment of NOD mice with $hSAgA_{mix}$, $cSAgA_{mix}$, $hp\text{-}peptide_{mix}$ or saline. (**B-I**) Expression of checkpoint molecules and cytokines on p79-reactive CD4+ T cells indicated as percentage of TIGIT+ (**B**), IL-2+ IL-10+ (**C**), IL-2+ (**D**), TNF-α+ (**E**), TNF-α+ IL-2+ (**F**), total IFN-γ+ (including cells co-expressing IL-10; **G**), KLRG1+ (**H**) and Lag3+ (**I**), all measured 3 days after the last of 23 treatments. Data in panels B-I show the mean ± SEM from 3-4 mice per group and two-way ANOVA/Tukey was used for statistical analysis.

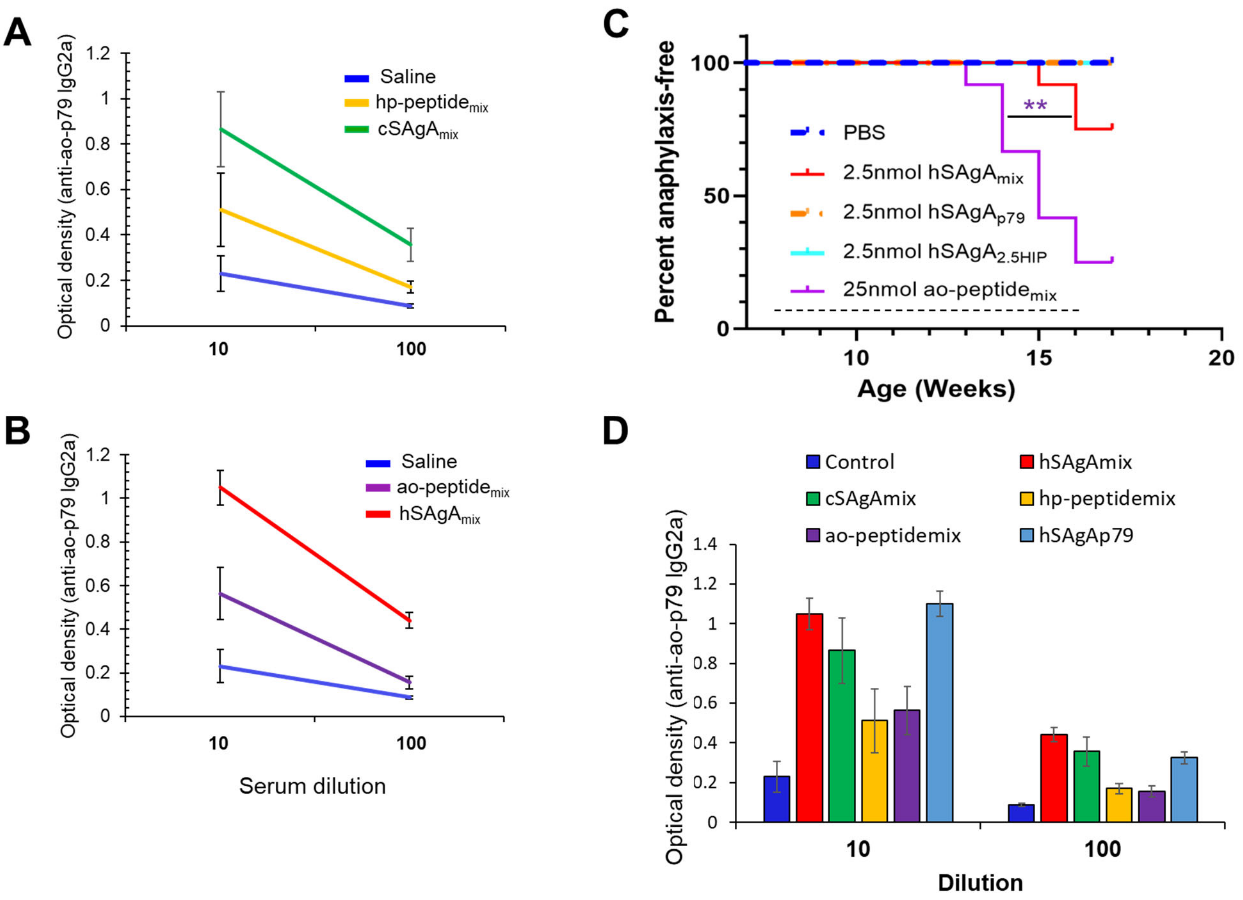


**Figure S5: Anaphylaxis incidence induced by a single hSAgA versus mix thereof (hSAgA$_{mix}$) and anti-p79 IgG2a response**. (**A**,**B**) Anti-p79 IgG2a levels were measured at the indicated serum dilutions following treatment with saline, hp-peptide$_{mix}$, cSAgA$_{mix}$, ao-peptide$_{mix}$ or hSAgA$_{mix}$. Data from cSAgAs/hp-peptides (A) and hSAgAs/ao-peptides (B) were separated for clarity. Data show the mean ± SEM from 4-5 biological replicates. (**C**) Incidence of anaphylaxis induced by a mix of the two hSAgAs (hSAgA$_{mix}$) or single SAgA carrying p79 or 2.5HIP at the indicated dose (n=12 per group). The dashed line indicates the weekly treatment period and log-rank test was applied for statistical analysis. The saline group of this figure was the same as saline group of Fig.1A as both experiments were done concurrently. (**D**) Anti-p79 IgG2a levels were measured at 1:10 and 1:100 serum dilutions following treatment with saline, hp-peptide$_{mix}$ or cSAgA$_{mix}$ (10 doses), or ao-peptide$_{mix}$, hSAgA$_{mix}$, or hSAgA$_{p79}$ (6 doses). The dose used was 2.5 nmol for cSAgA$_{mix}$ and hSAgA$_{mix}$, 5 nmol for hSAgA$_{p79}$, and 25 nmol for hp-peptide$_{mix}$ and ao-peptide$_{mix}$. The data represent the mean ± SEM from 4-5 biological replicates.

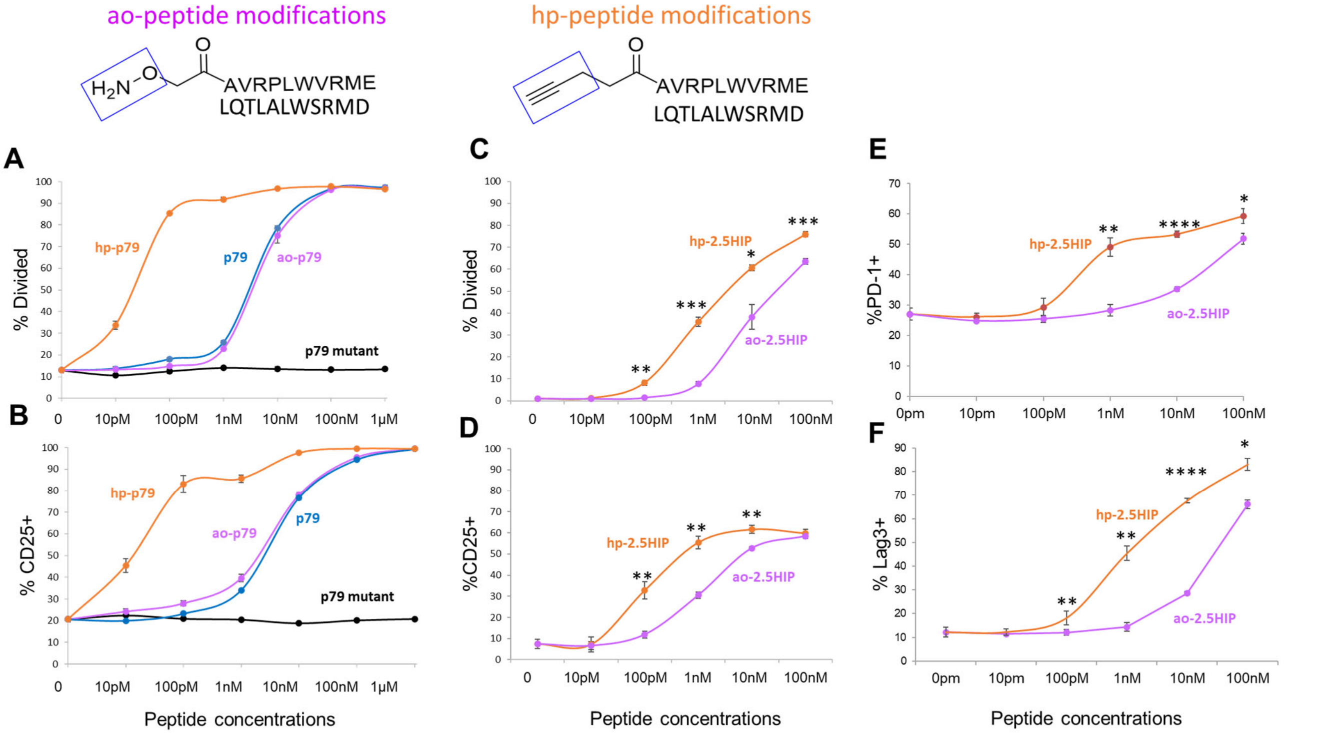


**Figure S6: Effect of peptide modifications on T cell stimulatory activity.** The ao-modified and hp-modified versions of p79 (A,B) or 2.5HIP (C,D) display a distinct potency in stimulating BDC2.5 CD4+ T cells. The response to p79 was also compared with unmodified peptide and mutant peptide (sequence AVPRLWVRME) (30). The T cell response was measured as proliferation (A,C) and CD25 upregulation (B,D), 3 days after culture of Carboxyfluorescein succinimidyl ester-labeled BDC2.5 splenocytes with titrated amounts of peptide. Upregulation of PD-1 (E) and Lag3 (F) in response to 2.5HIP is also shown. Data shows the mean ± SD from 3 technical replicates and non-parametric T-test was used for statistical analysis.
